# Transcatheter Edge-to-Edge Repair Increases Annular Forces in *In Vitro* Whole Heart Preparations

**DOI:** 10.64898/2026.03.04.709478

**Authors:** Collin E. Haese, Trace G. LaRue, Diego Guajardo, Campbell Harkness, William Hiesinger, Jan N. Fuhg, Tomasz A. Timek, Manuel K. Rausch

**Author notes:** These authors contributed equally to this work (Collin E. Haese, Trace G. LaRue).

## Abstract

**Background:** Tricuspid transcatheter edge-to-edge repair (TEER) can induce an acute annuloplasty effect. While this has a therapeutic benefit, the mechanisms driving the reduction in annular size remain unclear.

**Objectives:** We quantify the annular force induced by TEER *in vitro* in whole porcine heart preparations. We explore the impact of clipping different leaflet pairs on the TEER-induced annular forces.

**Methods:** We performed 49 interventions in 13 porcine hearts using a MitraClip XT. The clip was implanted between either the anterior-septal (AS), anterior-posterior (AP), or posterior-septal (SP) leaflet pairs. We also considered two-clip interventions between the combination of the AS-AP, AS-PS, or AP-PS leaflet pairs. For each intervention, we measured the right ventricular pressure, transvalvular flow rate, and force at eight locations around the annulus.

**Results:** TEER induced significant inward-pulling forces on the annulus. The maximum force was induced following an AS-PS two-clip intervention. A single AS clip induced the largest force among the one-clip interventions. Furthermore, the AP and AS-AP interventions induced the smallest annular forces.

**Conclusions:** The magnitude of the TEER-induced force depends on the intervention and number of clips implanted.

## 1. Introduction

Secondary tricuspid valve regurgitation (TR) is a disease that affects millions of individuals worldwide and is associated with increased mortality and morbidity [1]. Secondary, or functional, regurgitation is due to disease extrinsic to the valve and accounts for over 80% of all cases [2]. One of the driving mechanisms of secondary regurgitation is annular dilation. That is, the tricuspid annulus enlarges and impairs the natural coaptation of the leaflets and leads to gaps and leakage. Transcatheter edge-to-edge repair (TEER) has emerged as the most common percutaneous approach to treat TR. TEER addresses TR by approximating leaflets through the implantation of at least one “clip” to enhance coaptation and reduce regurgitation. TEER may also provide a therapeutic benefit by inducing acute annular remodeling resulting in a so-called “annuloplasty effect”. However, the mechanisms driving TEER-induced remodeling and the impact on outcomes remain unclear.

The TEER-induced annuloplasty effect has only recently gained attention as a potential therapeutic target. Several studies have reported on the reduction in annular dimensions following TEER [3, 4]. Importantly, von Stein et al. showed that a larger reduction in annular area was associated with better survival [5]. However, not all patients exhibit an annuloplasty effect following TEER. Russo et al. found that one-third of patients did not experience a reduction in annular dimensions [6]. Why some patients respond to TEER and others do not has not yet been explained and suggests an important knowledge gap.

Determining the mechanisms which cause the annuloplasty effect may fill a critical knowledge gap and help improve TEER outcomes. We submit that the annuloplasty effect is not well characterized in part due to the difficulty in studying TEER in a clinical setting. For example, the forces underlying the reduction in annular dimensions cannot be measured. In this study, we perform TEER *in vitro* in explanted porcine hearts under pathological pressures. We can measure the annular forces, right ventricular pressure, and transvalvular flow before and after intervention. Additionally, we can vary and compare different interventions within the same specimen, thereby overcoming several of the limitations of clinical studies. By quantifying the TEER-induced annular force, we aim to elucidate the mechanism driving the annuloplasty effect.

## 2. Methods

### 2.1. Experimental Setup

In this study, we used our previously-developed *in vitro* whole heart setup to subject the tricuspid valve to patho-physiologic conditions [7]. The setup is shown in Figure 1, and consisted of a whole porcine heart mounted in an acrylic chamber and submerged in 1 × PBS. A PBS reservoir was connected with 5/8” acetal tubing and a check valve to the right ventricle (RV) through the pulmonary artery and sealed with cable ties. A pressure transducer (5 psig PX409, Omega Engineering Inc., Stamford, CT) and electromagnetic flow meter (Model 501, Carolina Medical, East Bend, NC) were connected to this tubing to measure RV pressure and flow rate, respectively. Additionally, the reservoir was placed on a shelf of adjustable height which allowed for control of the static RV pressure. Furthermore, two high-resolution digital cameras (Imager CX-5, LaVision GmbH, Göttingen, Germany) captured atrial images through an acrylic viewing window which prevented surface distortions.

**Figure 1:**
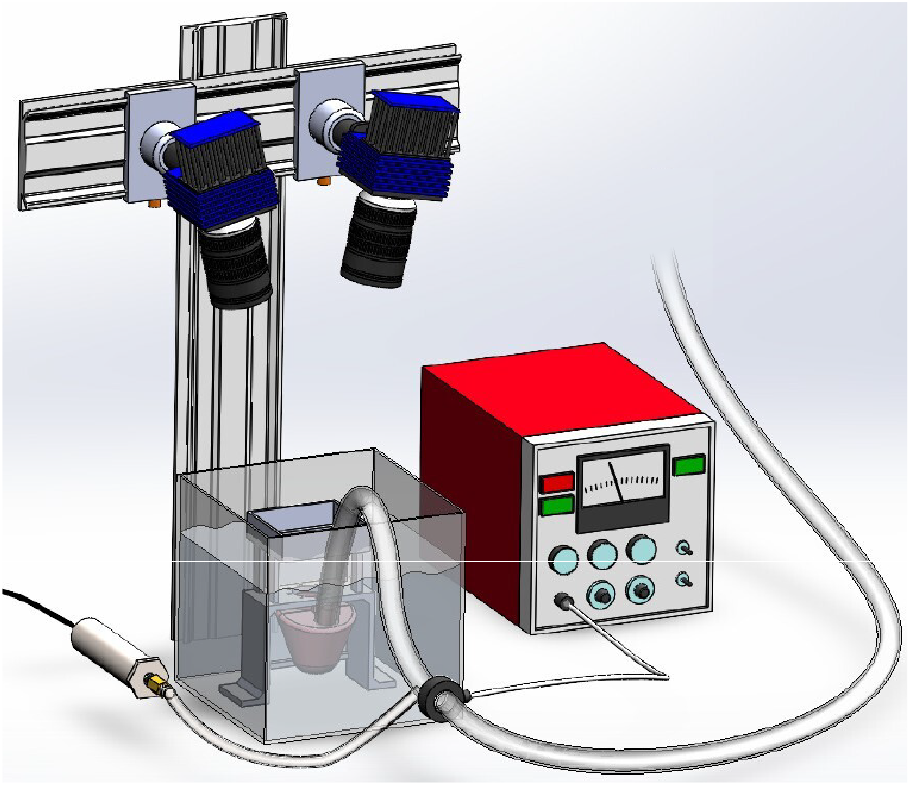
Our experimental, *in vitro* whole porcine heart setup. For each heart, we removed their atria, dilated their annulus and papillary muscles, mounted them in a fluid chamber, and connected the right ventricle to a static 1× PBS pressure reservoir. During testing, we pressurized the right ventricle and recorded right ventricular pressure, flow through the valve, and annular force at eight locations around the annulus.

### 2.2. Whole Porcine Heart Preparation and Mounting

We used thirteen (n=13) porcine hearts purchased from a local abattoir. All hearts were frozen and stored at −20 °C until testing [8]. Information about animal sex, age, and breed was not available. Figure 2 shows the preparation and mounting protocol used to prepare the hearts. Before testing, we removed both atria to expose the tricuspid annulus and enable attachment of the custom force transducers (Figure 2A). We also removed the pulmonary valve to allow unrestricted pressurization of the RV. We used a 3D-printed annular template based on an Edwards Classic size 30 annuloplasty device (Edwards Life-scienes, Irvine, CA) to mark eight, equally-spaced points around the tricuspid annulus (Figure 2B). At each of the eight points, we sutured one end of our custom force transducer to the tricuspid annulus and preserved the natural 3D height of the annulus (Figure 2C). See Section 2.3 for additional details on the force transducer. We used a template to mark the annulus to ensure a consistent placement of the force transducers across all hearts. We then attached the other end of the force transducer to our 3D-printed adjustable annular mount (Figure 2H, [7]). Lastly, we secured the annular mount to a stand fixed to the acrylic fluid chamber.

**Figure 2:**
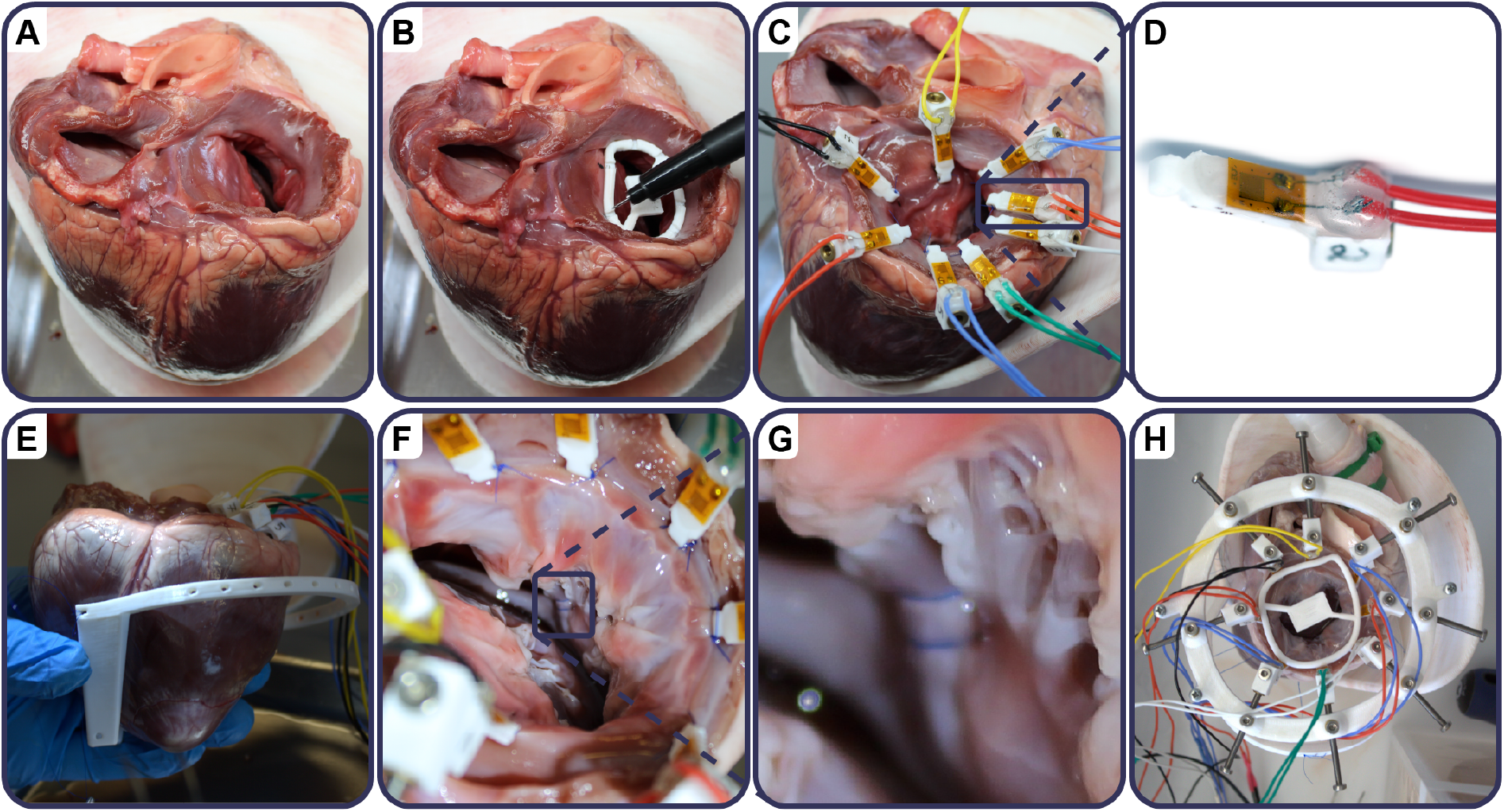
Whole porcine heart preparation and modeling of secondary tricuspid regurgitation. **A** First, we removed both atria and **B** uniformly marked the annulus at eight points. **C** At each mark, we sutured one end of our custom force transducers to the tricuspid annulus, i.e., at the junction of the tricuspid leaflets and myocardium. **D** A close-up view of one of the eight custom force transducers. **E** Next, we sutured a papillary muscle displacement ring to the interventricular septum. **F** We identified the papillary muscles on the free wall and **G** double-looped and externalized a suture around the papillary muscle and tied it through the papillary muscle displacement ring. We then placed a 15 mm spacer between the ring and suture. **H** Lastly, we connected all eight force transducers to the annular mounting ring and sized the annulus to an oversize annular template.

### 2.3. Custom Force Transducer

To measure the force around the tricuspid annulus, we designed and fabricated custom force transducers. To this end, we 3D printed a 12 mm× 4 mm ×1 mm pin in PLA. Next, we epoxied (LOCTITE EA M-31CL, Henkel, Düsseldorf, Germany) a linear strain gauge (SGD-3S/120-LY13, Omega Engineering Inc., Stamford, CT) to the surface of the pin. We soldered 28-gauge stranded-core wire to the strain gauge leads and sealed the gauge and solder with epoxy and hot melt adhesive to create a watertight connection. Wire leads were connected to a strain input module (NI-9236, National Instruments, Austin, TX) to measure voltage changes across the gauge. Finally, we calibrated the voltage change of the strain gauge while submerged against forces of known magnitude. We fabricated and calibrated eight such force transducers, one of which is shown in Figure 2D. Note, we refer to the force transducers as transducers or “pins” throughout the remainder of this study.

### 2.4. Model of Tricuspid Regurgitation

Since we acquired porcine hearts from a local abattoir, the tricuspid valves were functional and coapted even at large amounts of annular dilation alone [7]. To overcome this limitation of our previous study, we dilated the papillary muscles (PM) and tricuspid annulus in each heart to model secondary TR and induce leakage of the valve. That is, after preparing the heart and attaching the eight force transducers, we sutured a custom 3D-printed PM dilation ring (Figure 2E) to each side of the interventricular septum. We then rotated the ring to be parallel to the annular plane and sutured the bottom of the ring to the pericardium which prevented further rotation. Next, all the distinct papillary muscles located on the RV free wall were identified. We looped a suture twice around each PM, externalized the suture, and tied it taut through holes in the PM ring (Figure 2F-G). A 15 mm spacer was placed between the ring and suture to displace each PM by 15 mm [9]. Furthermore, we 3D printed oversize, pathologically dilated annular templates based on the Edwards Classic tricuspid annuloplasty rings (Edwards Life-sciences, Irvine, CA, [10, 11]). We created sizes 38-50 by inferring the annular area for each and asymmetrically dilating the size 36 profile to attain the target area and an ellipticity of one (Figure 2H, [12]).

### 2.5. Experimental Protocol

Our primary objective was to quantify the annular forces induced by TEER. Inspired by Vismara et al., we also aimed to maximize the number of TEER configurations tested in each heart to reduce the total number of specimens required [13]. Our experimental procedure for each heart is as follows. First, we placed our prepared and mounted heart in our acrylic fluid chamber and connected all tubes and wires. We placed our 1× PBS reservoir at the maximum height and matched the annulus to a size 38 oversize annuloplasty ring (Section 2.4). Next, the check valve was opened, the RV pressurized, and RV pressure recorded. We iteratively increased the annulus size using our adjustable annular mount to match the next largest oversize ring until the RV pressure reached 35 mmHg (Figure 2H). Once attained, the annulus was then locked at this shape and size. This configuration is referred to as the “control” configuration of the valve, which models an unrepaired tricuspid valve with secondary TR. We selected 35 mmHg to represent a typical pathological RV pressure seen in patients with secondary TR [14]. Again we used the check valve to simultaneously record the RV pressure, flow rate, atrial images, and annular force in all eight force transducers in the control configuration. This test was repeated three times, with each test lasting approximately 16 seconds. The average peak RV pressure measured across all three tests was selected as the “target pressure” for this heart. This completed the control experiment, and we then proceeded with TEER intervention of the valve.

We used the atrial images of the control valve to identify the leaflet pair with the largest coaptation gap. We then clipped across this leaflet pair with an Abbott XT MitraClip G4 (Abbott Structural Heart, Westfield, IN) on a custom delivery device, see Section 2.6. That is, with visual access we inserted the clip into the valve, captured both leaflets, and closed the clip as shown in Figure 3A-B. With the clip closed and still on the delivery device, we recorded the flow through the repaired valve. We then iteratively adjusted the location of the clip (by opening, recapturing leaflets, and closing) to minimize the flow through the valve. Once we attained the minimum observed flow, we again closed the clip and deployed it from the delivery device. Simultaneously, we adjusted the height of the 1× PBS reservoir until the RV pressure matched the target pressure from the control experiment. This post-TEER intervention configuration is referred to as the first “one-clip” configuration of the valve. We labeled this configuration as one of three interventions depending on the leaflet pair clipped: AS (anterior-septal), AP (anterior-posterior), or PS (posterior-septal). Again, we recorded the RV pressure, flow rate, atrial images, and annular force in this one-clip configuration and repeated this test three times.

**Figure 3:**
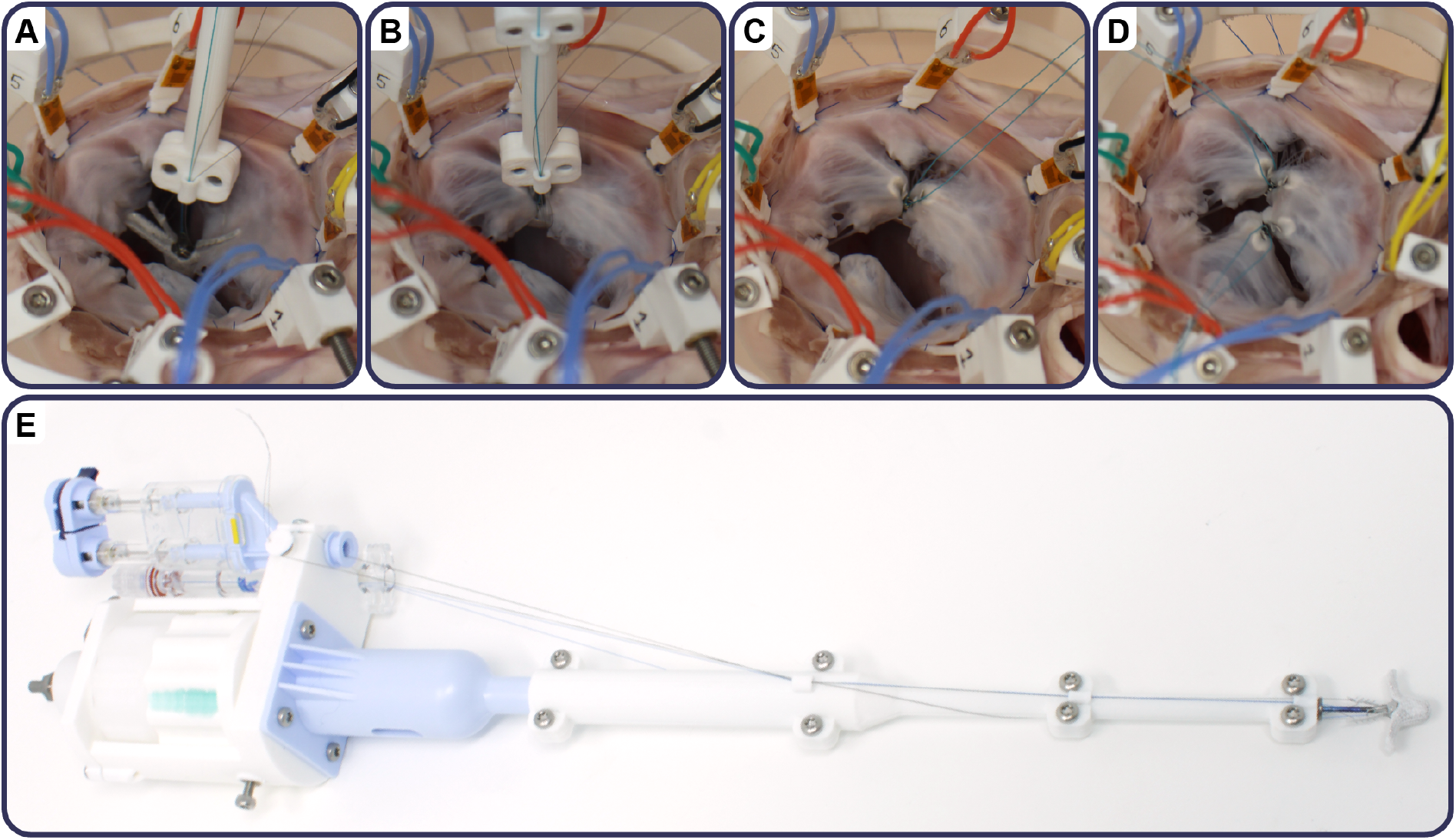
Our *in vitro* TEER procedure. **A** First, we insert the clip into the valve and grasp the first (septal) leaflet. **B** We grasp the second (anterior) leaflet and close the clip. Next, we test the flow through the valve and adjust the position of the clip between the two leaflets to minimize flow through the valve. **C**. After finding a suitable position, we deploy the clip by releasing it from the delivery device and test the one-clip configuration. **D** We follow the same procedure to deploy a second clip across a different leaflet pair, and test a two-clip configuration. **E** A detailed view of our custom clip delivery device.

Next, we used the atrial images of the one-clip valve to identify the next largest coaptation gap in a different leaflet pair. We clipped across this leaflet pair and adjusted the location to obtain a minimum flow through the valve. After deploying the second clip and adjusting the reservoir height to match the target pressure, we attained the first “two-clip” configuration of the valve. We labeled this configuration as one of three interventions depending on the the leaflet pairs clipped: AS-AP (AS and AP), AS-PS (AS and PS), or AP-PS (AP and PS). Again, we recorded the RV pressure, flow rate, atrial images, and annular force in this two-clip configuration and repeated this test three times. Lastly, we removed the first clip inserted to produce the next “one-clip” configuration. Again, we adjusted the reservoir height to match the target pressure and recorded the RV pressure, flow rate, atrial images, and annular force in this one-clip configuration three times. If time permitted, we would clip the remaining leaflet pair, test the second “two-clip” configuration, and then remove the second clip to test the third “one-clip” configuration. All testing was completed within 10 hours of thawing the heart.

At the conclusion of each experiment, we had recorded the RV pressure, flow rate, and annular force at eight locations for different combinations of clipped leaflet pairs. We never placed clips on the same leaflet pair, nor repeated leaflet pairs within the same heart. Additionally, we targeted a consistent pressure within a heart in order to remove the dependence of the annular force on the RV pressure alone.

### 2.6. Custom Clip Delivery Device

To perform TEER *in vitro*, we designed and fabricated a custom clip delivery device as shown in Figure 3E. The device was a modified version of the original Abbott device supplemented with 3D-printed components. Briefly, we made the delivery device handheld and ergonomic for use *in vitro* by shortening the catheter to 12”, removing the steering ability, and simplifying the deployment mechanism. We maintained the ability to independently raise/lower each gripper, lock/unlock the clip, open/close the clip, and deploy the clip. After each experiment, we removed all clips from the valve, cleaned them thoroughly, and loaded them back on to the delivery device for use in the next experiment. Altogether, these modifications allowed us to repeatedly perform TEER and reuse the clips across multiple experiments.

### 2.7. Data Analysis

We recorded all data using LabView 2023 Q4 (National Instruments, Austin, TX) and exported them as a .csv file. We then used MATLAB (Version R2024b, Mathworks, Natick, MA) to post-process these data. Therein, we zeroed the flow and force in each transducer at the start of the test. Next, we identified the starting and ending time points where all measurements reached quasi-equilibrium values. We then temporally averaged all measurements across this time period. For each transducer, we also computed the average change in force, or the TEER-induced force, by subtracting the average force in the transducer in the control configuration from the current intervention’s average force. We defined a positive force as one pushing radially outwards from the annulus, whereas a negative force is defined as pulling the annulus towards the center of the valve. Atrial images were recorded and exported as videos using DaVis 11 (LaVision GmbH, Göttingen, Germany).

### 2.8. Statistical Analysis

We report all values as means ± one standard error. We used the *afex* library in R (Version 4.1.2, The R Foundation, Vienna, Austria) to fit a linear mixed model to our data. This allowed us to test the dependence of annular forces on intervention, force transducer location, flow, and pressure, along with their interactions. All pairwise comparisons were performed using the “emmeans” library, also in R. We included each heart as a random effect in our model, and defined statistical significance as p*<*0.05.

## 3. Results

We performed 49 TEER interventions in 13 porcine hearts. We excluded two interventions due to 1) significant damage to a leaflet upon clip removal and 2) failure to reach the target pressure after clip deployment. We further excluded four tests from the data set due to significant noise or signal loss in at least one of the force transducers. For each intervention, we measured the RV pressure, flow rate through the valve, and force at eight locations around the tricuspid annulus. Figure 4 shows the raw data from one experiment. In this heart, one control, two one-clip, and one two-clip interventions were tested. First, we see that as the RV is pressurized at the beginning of the test, all measurements rapidly changed values. After the pressurization and closure of the valve, the measurements equilibrated to quasi-steady-state values for the remainder of the test. Additionally, we found that we were able to maintain nearly equivalent ventricular pressures across interventions while reducing flow, i.e. regurgitation, through the valve. In this heart, the two-clip intervention reduced flow more than either one-clip intervention. Furthermore, we found that the annular force changed between interventions and between annular locations. Specifically in this heart, force magnitudes were generally larger for the two-clip intervention. In summary, we were able to produce consistent quasistatic force, pressure, and flow measurements for each intervention tested.

**Figure 4:**
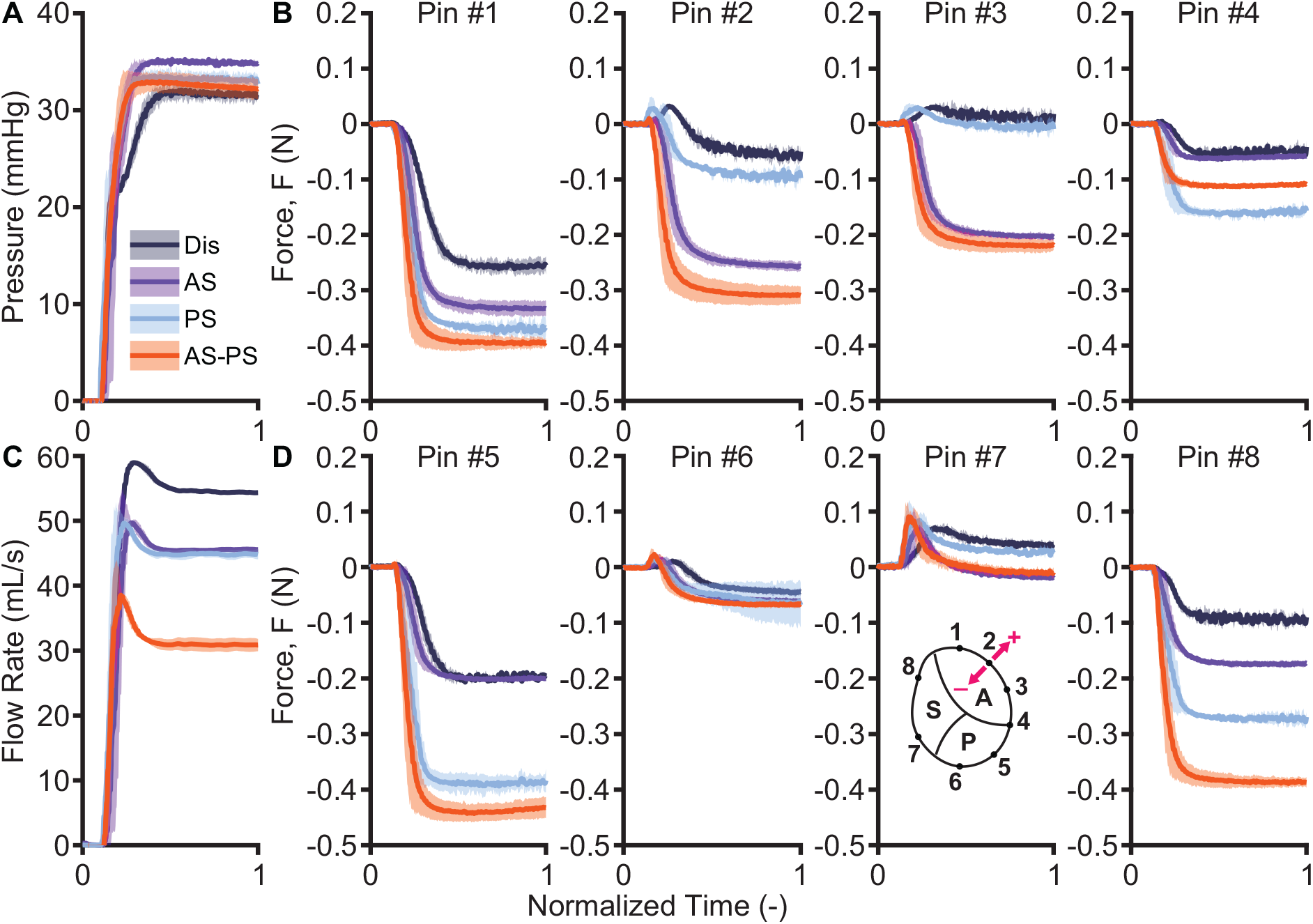
Sample raw data from three interventions tested in one porcine heart. We normalize and align all data measurements to a uniform time. For each intervention, we recorded measurements three times and plotted the average value bounded by one standard deviation. **A** Ventricular pressure throughout normalized testing time for each intervention. **B** Annular force in pins #1-4 throughout normalized testing time for each intervention. **C** Flow rate throughout normalized testing time for each intervention. **D** Annular force in pins #5-8 throughout normalized testing time for each intervention. Annular pin locations are shown inset under pin #7. A positive force is defined as pushing radially outwards from the annulus, whereas a negative force is defined as pulling the annulus towards the center of the valve.

### 3.1. TEER Reduces Transvalvular Flow

For each intervention tested within a given heart, our procedural target was to maintain the target ventricular pressure while minimizing the flow through the valve. Figure 5A shows the average RV pressure for each intervention tested. We maintained an average RV pressure of 36.04± 0.244 mmHg across all interventions. We found no statistical difference in pressure between any intervention position and the target pressure, except for the AS intervention (AS 37.22± 0.401 vs. CTL 35.85 ± 0.286 mmHg, p<0.001). Figure 5B shows the average flow rate across the valve for each intervention tested. We reduced the average flow rate from 52.69 ±0.448 mL/s in the control valves to 35.02 ±0.565 mL/s across all interventions. Each intervention significantly reduced flow with respect to control (AP p<0.01, all others p<0.001 vs. CTL). The AS-AP two-clip intervention showed the lowest average flow (29.70± 4.204 mL/s), followed by AS-PS, AS, SP, AP-PS, while the AP one-clip intervention showed the largest average flow (39.73±1.772 mL/s). Thus, we found that we were able to significantly reduce flow while maintaining equivalent ventricular pressures across hearts and interventions.

**Figure 5:**
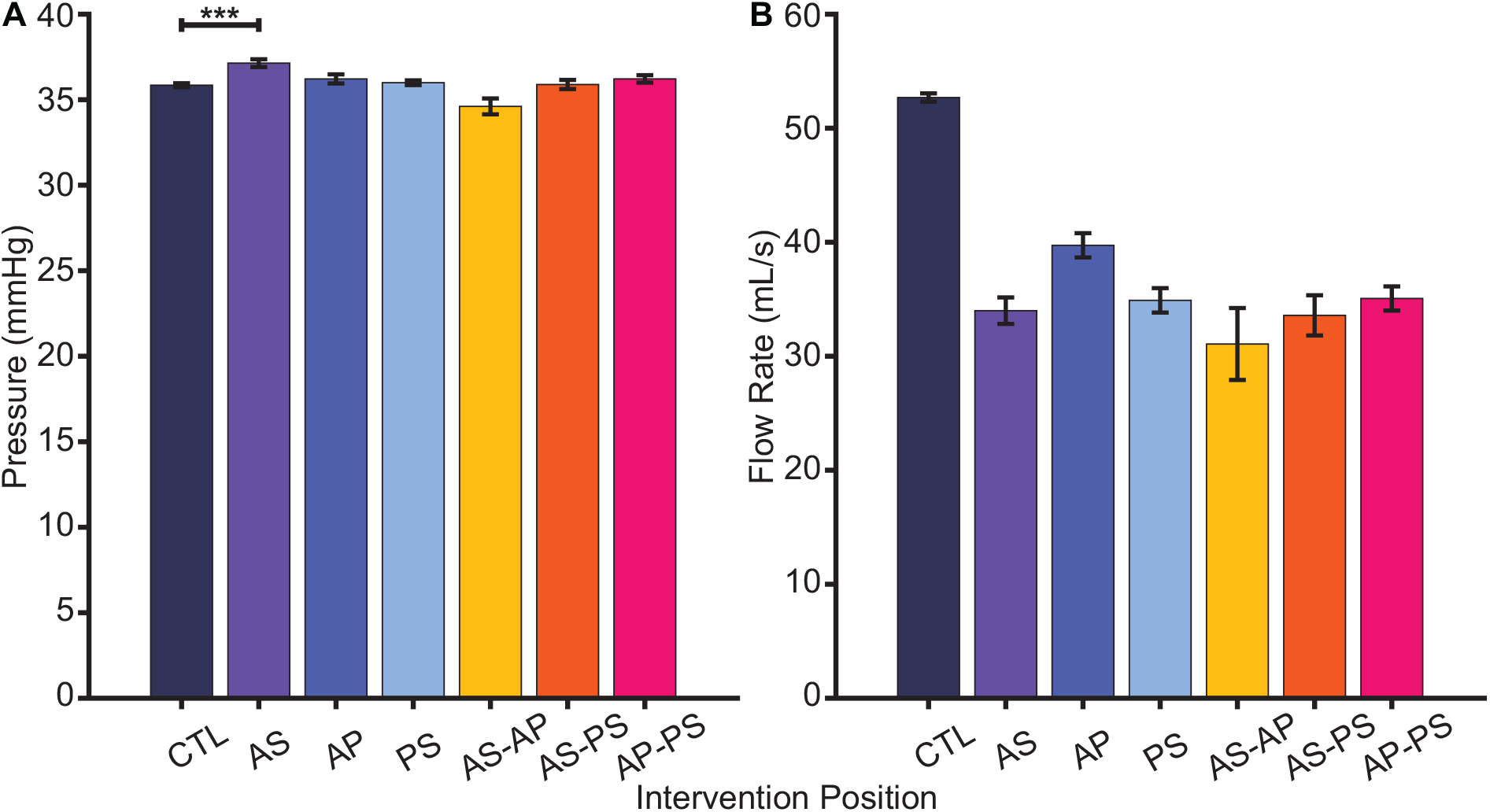
We held pressure fixed while reducing the flow rate across the valve. **A** Average right ventricular pressure for each intervention. **B** Average flow rate for each intervention, with each p<0.01 vs. control. Asterisks indicate statistical significance (* p<0.05, ** p<0.01, *** p<0.001).

### 3.2. Annular Forces Increase Following Intervention

To quantify and investigate the TEER-induced change in annular forces, we measured the force at eight locations around the tricuspid annulus before and after intervention. We found a statistically significant change in force following intervention (p<0.001). The largest change in force was observed following the AS-PS two-clip intervention (−0.118± 0.0174 N, p<0.001). A negative change in force represents a larger cinching or inward-pulling annular force induced by the clip. The other two-clip interventions also produced changes in force (AP-PS: −0.083± 0.0166 N, p<0.001; AS-AP: −0.039± 0.0175 N, p=0.067), but the AS-AP intervention failed to reach statistical significance. The largest change in force in the one-clip interventions was observed in the AS intervention (−0.055 ±0.0160 N, p<0.01), followed by the PS intervention (− 0.047± 0.0156 N, p=0.023). Interestingly, the AP intervention resulted in a slightly positive, but not statistically significant, change in annular force (0.002 ±0.0158 N, p=0.883). We also found a significant dependence of the change in force on the force transducer (pin) location (p<0.001). Pins #1, 2, 3, 5, 6, and 8 showed a significant negative change in force (pin #2 p<0.01, all others p<0.001), while pins #4 and 7 showed no statistically significant changes in force (p=1.000). Furthermore, we observed a significant statistical interaction between force transducer location (pin) and intervention (p<0.001). To further explore the intervention-specific change in force at individual pin locations, we next investigated this interaction between pin and intervention

### 3.3. Annular Forces Depend on Intervention

Figure 6 shows the TEER-induced force at each force transducer for the three one-clip interventions (AS, AP, PS). We found that an AS intervention induced significant inward-pulling forces along the anterior leaflet as measured in pins #2-3 and at the septal leaflet in pin #8, as shown in Figure 6A. Forces did not substantially change in any other transducers. For an AP intervention, we found significant inward-pulling force only along the anterior leaflet at pin #1 as shown in Figure 6B. There was significant outward-pushing force along the septal leaflet at pin #8, and the remainder of the pins (#2-7) did not show significantly changed forces. For a PS intervention position, we found significant inward-pulling forces on the anterior leaflet at pin #1, along the posterior leaflet at pin #5, and on the septal leaflet at pin #8 as shown in Figure 6C. All other pins showed non-significant changes in force, with pins #2-3 showing small outward-pushing forces. In summary, we found that the one-clip interventions induced annular force, the induced force tended to be inward-pulling on the annular segments connected to the clipped leaflets, while the untouched leaflet tended to show outward-pushing force.

**Figure 6:**
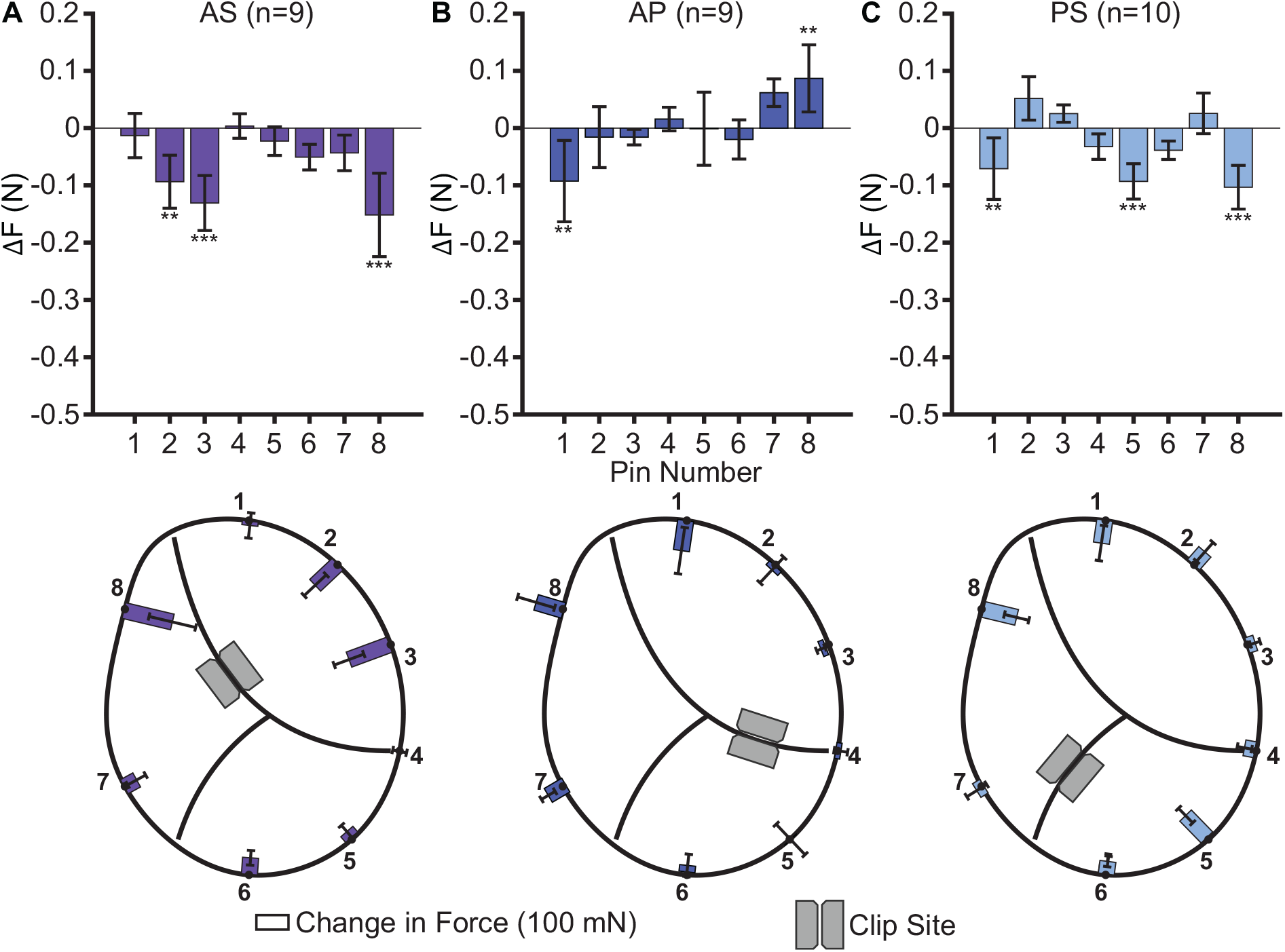
TEER-induced annular force for interventions with one clip. **A** One clip across the anterior-septal (AS) leaflet pair induces significant inward-pulling forces at pins #2, 3, and 8. **B** One clip across the anterior-posterior (AP) leaflet pair induces significant inward-pulling force only at pin #1, and induces significant outward-pushing force at pin #8. **C** One clip across the posterior-septal (PS) leaflet pair induces significant inward-pulling forces at pins #1, 5, and 8. Asterisks indicate statistical significance (* p<0.05, ** p<0.01, *** p<0.001) with respect to control. A positive force is defined as pushing radially outwards from the annulus, whereas a negative force is defined as pulling the annulus towards the center of the valve. Sample size for each intervention given as n=.

We also quantified the annular force for the two-clip interventions. Figure 7 shows the TEER-induced force at each force transducer following intervention with two clips. We found that an AS-AP intervention induced significant inward-pulling forces only at the anterior leaflet in pins #2 and 3 as shown in Figure 7A. Forces were not significantly changed in all other pins. For the AS-PS intervention, we found significant inward-pulling forces along all three leaflets at pins #2, 3, 5, 6, and 8. The remaining pins showed no statistical change as shown in Figure 7B. Finally, for a AP-PS intervention we again found significant inward-pulling forces along all three leaflets at pins #1, 2, 5, 6, and 8 as shown in Figure 7C. All other pins showed non-significant changes in force. Overall, we found that two-clip interventions tended to induced inward-pulling annular forces around the annulus at all pins except pins #4 and 7.

**Figure 7:**
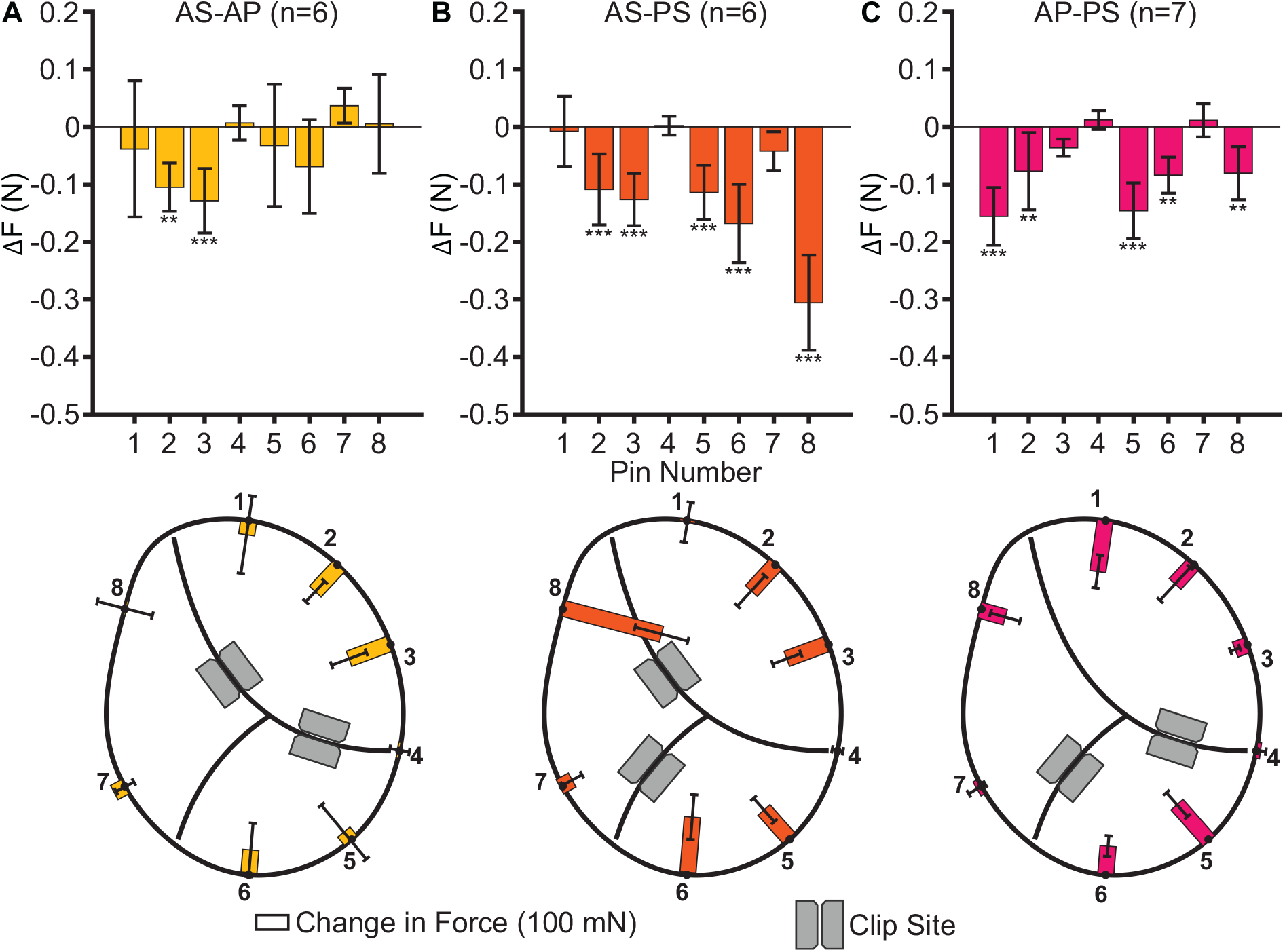
TEER-induced annular force for interventions with two clips. **A** Two clips across the anterior-septal and anterior-posterior (AS-AP) leaflet pairs induces significant inward-pulling forces at pins #2 and 3. **B** Two clips across the anterior-septal and posterior-septal (AS-PS) leaflet pairs induce significant inward-pulling forces at pins #2, 3, 5, 6, and 8. **C** Two clips across the anterior-posterior and posterior-septal (AP-PS) leaflet pairs induces forces at pins #1, 2, 5, 6, and 8. Asterisks indicate statistical significance (* p<0.05, ** p<0.01, *** p<0.001) with respect to control. A positive force is defined as pushing radially outwards from the annulus, whereas a negative force is defined as pulling the annulus towards the center of the valve. Sample size for each intervention given as n=.

## 4. Discussion

In this study, we used our *in vitro* whole porcine heart preparations to quantify TEER-induced forces around the tricuspid annulus. We performed 49 interventions with a MitraClip XT in 13 hearts. We varied the number of clips inserted (one or two) and the leaflet pairs clipped (AS, AP, PS). We measured the force at eight annular locations using custom force transducers and computed the TEER-induced force as the difference between the mean pre- and post-intervention force. For each test, we also measured the transvalvular flow while keeping the RV pressure consistent within a specimen. We found minor differences in pressure across interventions, but were able to significantly reduce flow through the valve. We also found a significant change in annular force, and found that this change depended on the intervention and force transducer location.

We found that TEER induced inward-pulling annular forces. This increased force tended to act on the segments of the leaflet pair which was clipped. For example, in the AS intervention only the forces on the anterior and septal leaflets significantly increased as seen in pins #2, 3, and 8. Furthermore, adding a second clip appeared to superimpose the forces induced by each clip individually. In the AS-PS intervention, for example, the force on all three leaflets significantly increased as seen in pins #2, 3, 5, 6, and 8. We also found that the AS-PS intervention, the so-called “clover technique”, induced the most annular force. On the other hand, the AP and AS-AP interventions tended to induce the lowest forces. This supports current practice in which the AP leaflet pair is rarely clipped.

To the best of our knowledge, this is the first study to quantify TEER-induced force in the tricuspid annulus *in vitro* in whole hearts. We chose to quantify the annular force as it is a likely mechanism driving the annuloplasty effect. While there are no other *in vivo* or *in vitro* reports of TEER-induced force, our results align with many observational findings. Widmann et al. suggested that TEER applies traction to the leaflets which leads to annular remodeling [4]. While they were unable to measure this traction, this study confirms that TEER does apply force to the clipped leaflets. Our findings further align with recent reports by von Stein et al. that associated the clover strategy (AS-PS intervention) with a larger degree of annuloplasty [5]. However, they were comparing to a “zipping” strategy were two or more clips are placed between the same leaflet pair. While we did not test two clips between the same leaflet pair, we did observe that adding additional clips between different leaflets appeared to superimpose the force contribution from each clip alone. We believe that adding a second clip between the same leaflet pair would do little to further increase the TEER-induced annular force as the first clip already applies the traction on the valvular apparatus. Additionally, Russo et al. observed that implanting two clips was associated with a larger reduction in annular area than one clip [6]. Our data supports this finding as the two-clip interventions were found to induce larger annular forces. Importantly, we showed *in silico* that TEER induces significant annular forces [15]. In this prior study, we showed that the observed annuloplasty effect correlates with the magnitude of the TEER-induced annular forces, that the induced annular forces depend on clip orientation, clip size, and clip site (leaflet pair). Our current study corroborates and supports the findings of this prior study.

Importantly, our results offer several translational take-aways. First, we quantify the TEER-induced force for six different interventions. We found that the induced force increases following TEER. Furthermore, our results demonstrate that the magnitude of the induced force varies between interventions and specimens. This reinforces the need for subject-specific guidance when performing TEER. Second, we found that the annular forces tended to increase on the leaflet pair clipped. That is, the clip can be used to direct force to adjacent segments of the annulus. This can be used to target compliant sections of the annulus and RV free wall to maximize the TEER-induced annuloplasty effect. Lastly, we found that the two-clip interventions on average induced more force than the one-clip interventions. This finding supports the use of multiple clips when performing TEER [16]. However, we note that not all two-clip interventions induced more force than one-clip counterparts, again highlighting the need for a subject-specific TEER strategy.

### 4.1. Limitations and Future Work

This study is an *in vitro* investigation of the effects of TEER on the annular force in the tricuspid valve. We performed TEER with clinical devices in real hearts. However, these hearts were explanted porcine hearts obtained from a local abattoir. As such, they were functional and their leaflets and ventricles lacked the remodeling characteristics of patients with tricuspid regurgitation [17, 18]. This likely resulted in more compliant and thinner structures, which may have led to an underestimation of annular forces. In the future, we will aim to conduct similar experiments in functionally remodeled hearts. Furthermore, the explanted hearts lacked contractility and the dynamic motion of the tricuspid annulus present in a beating heart. However, we were required to fix the annular shape in order to measure the annular forces. We used oversize annular templates based on commercially available annuloplasty rings to enforce physiologic, end-systolic shapes. This assumption likely impacted the resulting leaflet kinematics and annular forces, and precluded the observation of an annuloplasty effect. In future studies, we will instead opt to allow annular displacement and measure the annuloplasty effect directly. Lastly, we aimed to standardize the location of the force transducers around the annulus by using a standard template to uniformly mark the annulus. This could not fully account for morphological differences across each valve, and therefore some of the transducers may have been in slightly different anatomic positions. Additionally, pins #4 and 7 were often located near the anterior-posterior and posterior-septal commissure, respectively. The leaflets tend to fold at the commissures and transmit little force to the annulus. This likely explains why only these two transducers did not see a statistically significant change in force across interventions. We can instead aim to attach the force transducers based on anatomical markers in future studies.

## 5. Conclusion

We found that TEER induces significant annular forces *in vitro* in whole porcine hearts. The TEER-induced force depends on the intervention, i.e., on the leaflets that are clipped. Two-clip interventions generally increase annular force more than one-clip interventions. Additionally, clipping across the anterior-posterior leaflets appears to induce the smallest amount of annular force. Forces tend to increase in the segments of the annulus along the clipped leaflets. Furthermore, forces appear to superimpose and add when additional clips are implanted. Leveraging this clip-induced force may help to maximize the amount of annular remodeling. In summary, TEER induces significant annular forces across the clipped leaflets which likely drives the amount of TEER-induced annuloplasty.

## 6. Acknowledgments

This research was supported by funding from the National Heart, Lung, and Blood Institute of the National Institutes of Health through award numbers 1R01HL165251 and 1R21HL161832 to M.K.R. and T.A.T., and 1F31HL178280 to C.E.H. We also thank Margot Vandenheede for her insights and suture skills while performing some of the experiments.

## 8. Data Availability Statement

All raw data files, images, and statistical codes are openly available through the Texas Data Repository.

